# Evidence of a role for radiation-induced lysosomal damage in non-targeted effects: an experimental and theoretical analysis

**DOI:** 10.1101/024661

**Authors:** Scott Bright, Alexander G. Fletcher, David Fell, Munira A. Kadhim

**Affiliations:** Department of Biological and Medical Sciences, Oxford Brookes University, Oxford, OX3 0BP, UK; Mathematical Institute, University of Oxford, Andrew Wiles Building, Radcliffe Observatory Quarter, Woodstock Road, Oxford OX2 6GG, UK

## Abstract

A well-known DNA-damaging agent and carcinogen, ionizing radiation (IR) can also exert detrimental effects in cells not directly exposed to it, through “non-targeted effects” (NTE). Whilst NTE are known to contribute to radiation-induced damage, their mechanism of induction and propagation remains incompletely understood. To investigate the possible role of lysosomes, key subcellular organelles, in NTE we used acridine orange uptake and relocation methods to monitor lysosomal permeability in irradiated and bystander human fibroblasts. As a potential mediator of lysosomal changes, oxidative stress was measured using the H_2_DCFDA assay for total reactive oxygen species (ROS). IR was found to induce significant lysosomal permeability in the first hour post irradiation, with reduced permeability persisting up to 24 hours. This occurred in conjunction with an increase in ROS in directly irradiated cells, in contrast with a decrease in ROS in bystander cells. Based on these observations we constructed a simple mathematical model of ROS-induced lysosomal damage, based on a bistable mechanism where a sufficiently strong IR insult can shift a cell from a ‘low ROS, high lysosome’ to a ‘high ROS, low lysosome’ state. This has profound cellular implications in radiation response and advances our understanding for the sub-cellular involvement in non-targeted effects.

## Introduction

Ionizing radiation (IR) is a continual presence in our environment from natural and man-made sources *(1)*. IR interacts alters the structure and, subsequently, function of biological cells *(2)*. To date, radiobiological studies have to a large extent focused on the direct interaction of IR with nuclear DNA. It is widely accepted that IR is able to damage DNA, inducing mutations, chromosomal rearrangements, the formation of micronuclei and cell death *(2, 3)*. However, over the last 20 years the paradigm of radiation biology has undergone a transition from one of *targets* (biological effects induced by IR interacting with a specific target, namely, nuclear DNA) to one which incorporates *non-targeted effects* (NTE) *(4-8)*. Non-targeted effects are radiation-like effects observed in cells that have not been irradiated but which are the progeny of, or have communicated with, cells that have been irradiated; these two phenomena are termed genomic instability (GI) and bystander effects (BE), respectively *(5, 6, 9)*. While BE and GI have been well documented, their precise underlying mechanisms remain poorly understood, though it is likely that IR-induced sub-cellular signalling is involved through mechanisms such as inflammation *(10-12)*.

One proposed mechanism of IR-induced NTE involves reactive oxygen species (ROS) *(12-14)*. ROS are chemical species that carry an unpaired electron and are therefore highly reactive; they are produced in abundance following IR *(2)*. Although ROS are essential for cellular homeostasis through various signalling pathways such as cell migration and immune processing *(15, 16)*, an excess of ROS can result in cell damage and cell death *(16)*. ROS can also alter the structure and function of a substrate, leading to the formation of radicals. Such reactions can cause significant biological damage and are responsible for much of the IR-induced damage that arises in the cell *(2)*. It is therefore important to understand potential reactions and the downstream effects for cells in which they occur.

In the present study we evaluate the ability of IR to induce lysosomal permeability and explore a potential role for ROS in mediating this process (Figure 1). Damage to lysosomes has previously been demonstrated using chemical compounds such as sphingosine *(17, 18)*, the effect of which was cell death *(19-21)*. Allison and Paton [19] also demonstrated that lysosomal breakdown caused by lysomotrophic chemicals resulted in the appearance of radiation-like chromosomal aberrations. It was proposed that enzymes held within the lysosome, in particular DNase IIα, were responsible for these effects. Lysosomes also contain many enzymes, such as acid sphingomylinase and RNases, that can be involved in signalling *(22-24)*. More recent studies have investigated the mechanism of lysosomal breakdown and indicated that lysosomes are susceptible to ROS-induced damage, particularly through Fenton chemistry *(25, 26)*. Such reactions are prevalent due to the high iron concentration within the lysosome *(27)*. Owing to the potential cytotoxic and sub-lethal release of lysosomal enzymes, and the ability of IR to induce ROS to excess in the intracellular environment through physical reactions with water *(28, 29)*, the effect of IR and excess ROS on the lysosomal membrane is of particular interest.

**Figure 1:**
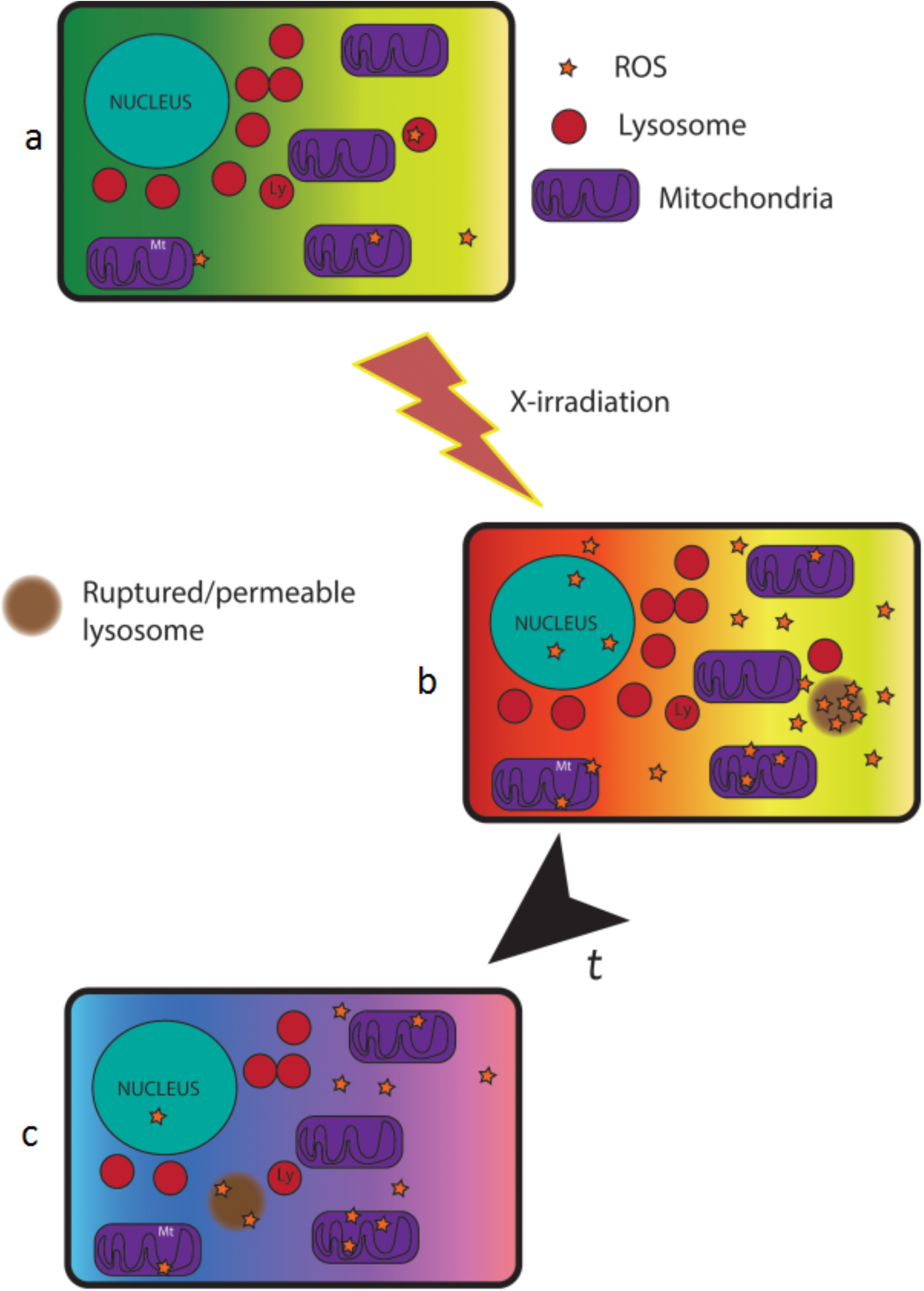
Under normal conditions, intracellular numbers of lysosomes and ROS are stable, resulting in no harm to the cell (a). However, radiation has the ability to damage cells, directly by inducing direct DNA damage or indirectly through the increased production of ROS molecules. Cellular damage could also occur through alterations in sub-cellular organelles, the total cellular damage may involve all of these processes (b). There is then a period of recovery (t) however processes are still ongoing in the cell as a result of the IR exposure (c) such as increased ROS that can still damage the cell.

Here we explore the lysosomal response to diagnostic and therapeutic doses of IR in the form of X-rays, with particular attention on ROS and lysosomal interactions, in the primary human fibroblast cell line HF19. We consider both directly irradiated cells and bystander populations (cells that received media from irradiated cells 4 hours post irradiation). We measure DNA damage using the comet assay, while lysosomal permeability and number are assessed over a 24-hour period in directly irradiated and bystander groups, in conjunction with oxidative stress. To further investigate the possible mechanism underlying the temporal dynamics of ROS and lysosomal permeability following IR exposure, we develop and analyse an ordinary differential equation (ODE) model of the intracellular interactions between ROS and lysosomes.

## Materials and methods

### Cell culture and irradiation

HF19 cells, a primary human fibroblast cell line *(30)*, were routinely sub-cultured in Minimum Essential Medium (MEM) supplemented with 2 mM L-glutamine, 50,000 units Penicillin, 0.1 mg/ml Streptomycin and 75ml foetal bovine sera (FBS). Sub-culture took place between 48 and 72 hours following the previous sub-culture. HF19 cells typically have a population doubling time of 36 hours, cell cycle distribution was not measured at point of measurements. Adherent cells were detached using 0.25% trypsin and propagated. Cell irradiations were performed at the Gray Institute for Radiation Oncology & Biology, Department of Oncology, University of Oxford, using an MXR321 X-ray machine operating at 250 kV. All irradiations were carried out in tissue culture flasks or tissue culture plates. Doses of 0.1 Gy, relevant to diagnostic procedures, or 2 Gy, relevant to therapeutic procedures, were used at dose rates of 0.59 and 0.58 Gy/Min respectively.

### Comet assay

The comet assay was performed in alkaline conditions to measure total DNA damage *(31, 32)*. Standard microscope slides were dipped in 1% normal melting point agarose (NMPA). The excess was wiped from the back and the slides were air dried overnight, they were then stored in a microscope box. Prior to cell harvesting, alkaline lysis buffer (2.5 M NaCl, 100 mM EDTA (pH 8.0), 10 mM Tris-HCl (pH 7.6), 1% Triton X-100, pH to 10), alkaline electrophoresis buffer (0.3 M NaOH, 1 mM EDTA, pH 13) and neutralization buffer (0.4 M Tris-HCl, pH 7.5) were prepared and chilled to 4 °C.

On the day of harvest, the cells were trypsinized and collected in fresh media, and cell counts were performed. Aliquots of 20,000 cells per group were put into 1.5 ml eppendorf tubes and placed immediately on ice. When all samples had been collected, the cell suspension was mixed with 200 μL of 1% low melting point agarose (LMPA). The NMPA pre-coated slides were placed on an ice-chilled metal plate and 200 μL of the LMPA cell suspension mixture was pipetted on to the chilled slide; a 22 x 50 mm glass cover slip was placed on top. Each slide was left for 5 minutes to allow complete setting, after which the cover slips were removed and the slides put into alkaline lysis buffer; Triton-X and DMSO (1%) were added 30 minutes prior. Lysis was carried out at 4 °C in the dark. The lysis time ranged from 16 to 20 hours but was consistent within time points.

After lysis treatment, the slides were equilibrated in the alkaline electrophoresis buffer in a horizontal tank. Subsequently, the slides underwent electrophoresis for 30 minutes in the dark at 19 V. The slides were then removed from the tank and neutralized with three 10 minute washes with neutralization buffer. Any remaining buffer was removed with 4 washes of dH_2_O. Slides were stained with 1:10,000 dilution of SYBR Gold in dH_2_O for 5 minutes in the dark. Finally, the slides were dried at 25°C overnight and analysed on a Nikon eclipse 600 microscope at 200x magnification using Komet 5.5 Image Analysis Software (Kinetic Imaging Technology/Andor, Germany).

### Lysosomal permeability

Lysosomal permeability was measured using acridine orange (AO) uptake and relocation methods (Figure 2) *(33-36)*. HF19 cells were grown on 13 mm diameter number 0 cover slips in 35 mm petri dishes at a density of 1.6 x 10^5^ cells per dish. All cells were stained for 15 minutes with 5 μg/ml of AO under standard culture conditions. The method is based upon the metachromatic properties of AO, where at high concentrations it fluoresces red and at lower concentrations it fluoresces green. AO freely diffuses across the cell membrane, and is sequestered into lysosomes at high concentrations. The residual dye remains in the cytoplasm at lower concentrations where it fluoresces green.

**Figure 2:**
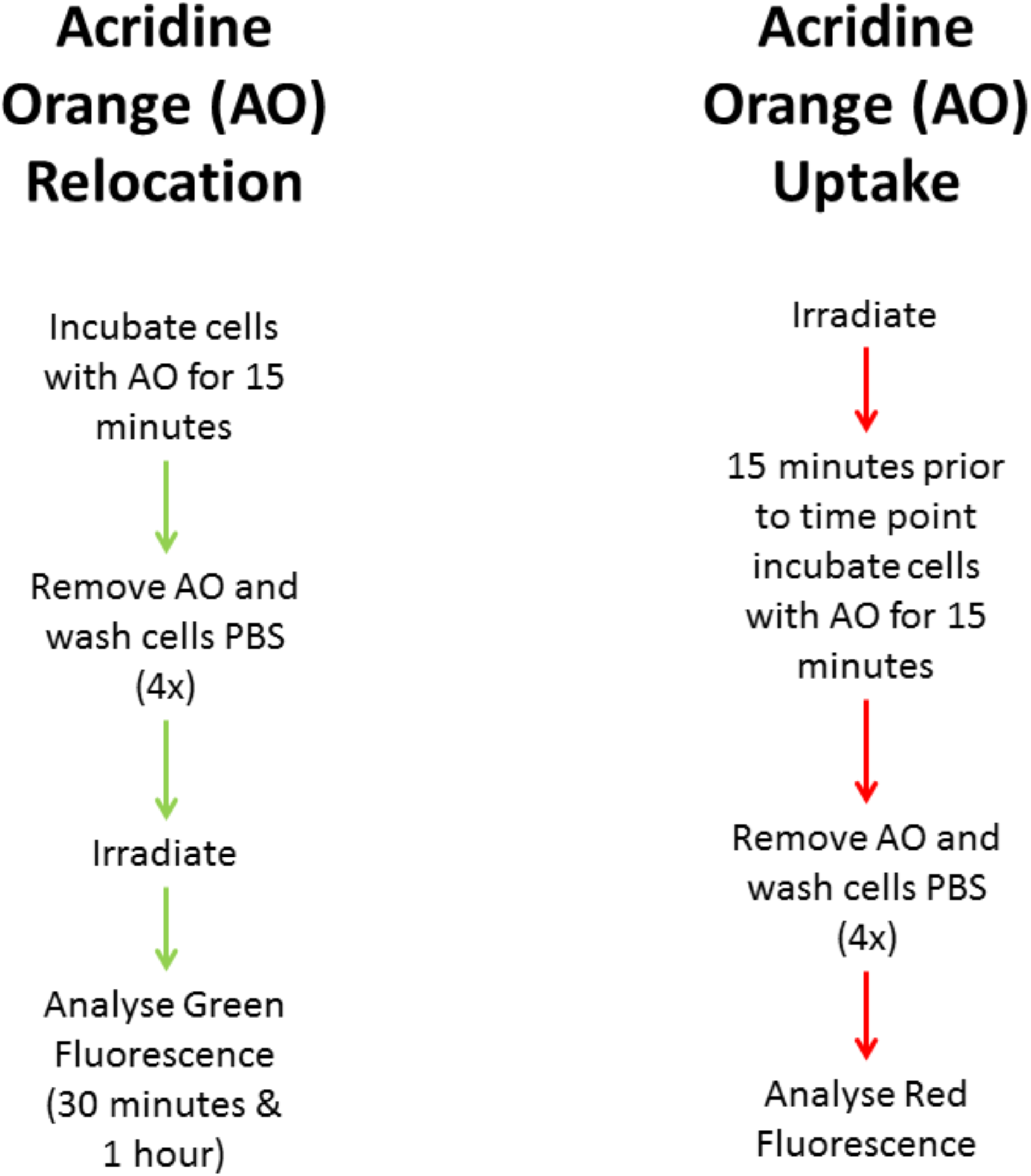
Acridine orange uptake and relocation methods were used depending on the time to be analysed. For time points up to 1 hour post irradiation cells were pre-loaded with AO and analysed for green fluorescence. Cells that were analysed further than 1 hour post irradiation were analysed according to AO uptake method, in this instance cells were loaded with AO 15 minutes prior to scheduled analysis time point.

### Acridine orange relocation

AO relocation requires pre-loading of the cells (i.e. the cells are stained prior to radiation exposure). It is then possible to monitor an increase in green fluorescence as lysosomes (red) become more permeable and leak AO into the cell cytoplasm (green). Experimental time points measured using AO relocation methods were 30 minutes and 1 hour post irradiation. In this instance cells were loaded with 5 μg/ml AO for 15 minutes under standard culture conditions. Following staining, cells were washed twice with PBS and fresh media was applied. They were subsequently irradiated and then prepared for imaging at the appropriate time point.

### Acridine orange uptake

AO uptake focuses on the principle that at delayed time points following radiation exposure, rupture will have already occurred, and therefore instead of monitoring the transition from red to green it uses total red fluorescence as an indicator of permeability. This method was employed for time points from 4 hours post irradiation. Fifteen minutes prior to the time point cells were loaded with 5 μg/ml AO for 15 minutes. Individual cover slips were washed 4 times with PBS and prepared for imaging.

### Imaging lysosomal membrane permeability

As stated previously, cells were grown on ethanol-sterilised cover slips in 35 mm petri dishes for at least 24 hours prior to testing. After cell loading with AO, individual cover slips were washed 4 times in warmed PBS (37 °C), and then mounted in warmed PBS onto a microscope slide. The cover slips were slightly raised from the slide surface by two parallel pieces of tape running horizontally across the slide approximately 10 mm apart. The cells were imaged using a 488 nm argon laser with a 505-550 nm BP filter for green fluorescence and a 615 nm LP filter for red fluorescence on a Zeiss LSM 510 meta upright laser scanning confocal microscope.

### ROS measurement

The general ROS marker CM-H_2_DCFDA (2’,7’dichlorodihydrofluorescein diacetate) was used to measure intracellular ROS levels in HF19 post irradiation. HF19 cells were seeded into black 96-well plates and measured using the Tecan Infinite F200 pro plate reader (Ex:488/Em:535). Cells were seeded at 7000 cells/well at 20-24 hours prior to measurement. The dye was prepared immediately before use. The dye was made into a stock solution of 1 mM in pure ethanol and was subsequently diluted in warmed PBS (37 °C) and used at a final concentration of 5 μM.

Media was removed from the wells to be tested and replaced with the PBS, containing CM-H_2_DCFDA. Cells were incubated for 30 minutes under standard culture conditions. After 30 minutes the PBS containing dye was removed and replaced with fresh media. Cells were allowed a 30-minute recovery period under standard culture conditions and then measured using the plate reader with filters of Ex:488/Em:525.

### Media transfer

In order to investigate radiation induced bystander effects, media from irradiated cells was transferred onto HF19 cells, the cells that received this media were called bystander cells *(37)*. Bystander cells were grown in identical fashion to directly irradiated cells. HF19 cells were irradiated with 0, 0.1 or 2 Gy X-rays and returned to standard culture conditions for 4 hours at which time the media was aspirated and filtered through a 0.22 μm filter that was pre-treated with 1% BSA in PBS. The media from the recipient bystander population was removed and discarded, the bystander cells were washed once with pre-warmed PBS and the filtered media was applied.

### Statistics

Comet assay results are representative of 200 cells across two biological replicates; these data were statistically analysed using the Kruskal Wallis test. Lysosomal data are representative of 50 cells scored across 3 biological replicates; these data were statistically analysed using the Mann Whitney test. Oxidative stress was analysed in 16 wells of a 96-well plate in 2 biological replicates; these data were statistically analysed using a 2 tailed t-test.

### Mathematical modelling

To complement our experimental study, we develop a simple mathematical model of the temporal dynamics in the numbers of lysosomes and ROS in a cell prior to, during and following IR. Our aim is to use the model to address hypothesised effect(s) of IR on ROS dynamics, and hypothesised interaction(s) between ROS and lysosomes. For simplicity, we restrict our focus to directly irradiated cells.

In our model, we assume that the number of each species may be treated as a continuous quantity and that a mean-field approximation is valid, thus neglecting stochastic fluctuations. We also neglect intracellular localisation of lysosomes and ROS, thus treating the cell as spatially homogeneous. We therefore use an ordinary differential equation (ODE) approach to model the abundance of each species. As a first approximation, we neglect longer-term dynamics in these quantities over the course of the cell cycle; in particular, we do not take into account the gradual doubling in lysosomal numbers that must occur in each cell prior to division. This form of modelling approach has been successfully used to shed light on the design principles and dynamics of molecular regulatory networks and such as the cell cycle oscillator and the phosphorylation cascade controlling bacterial chemotaxis, as well as other intracellular interactions; see, for example, Tyson *et al.* 2003 *(38)*.

We consider the following processes to affect levels of lysosomes and ROS in the cell, all of which we assume to be governed by mass action kinetics: ROS are generated constitutively at a constant rate *k*_1_; lysosomes are generated constitutively at a constant rate *k*_2_; ROS decay constitutively at a constant rate *k*_3_; lysosomes decay constitutively at a constant rate *k*_4_; and IR instantaneously elevates the rate of generation of ROS to *ck*_1_;, where the parameter *c* is related to the dose. In addition, we consider that ROS ‘bind to’ lysosomes and cause them to rupture, at a constant rate *k*_5_;. We suppose that this reaction *m* requires ROS per lysosome, and that the resulting lysosomal rupture generates a net increase of *n* ROS. The temporal evolution in the numbers of ROS and lysosomes, denoted by *R*(*t*) and *L*(*t*) respectively, is thus governed by the following ODE system

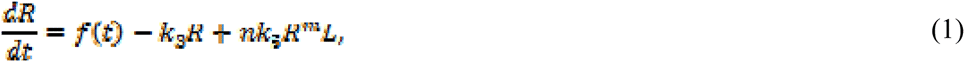

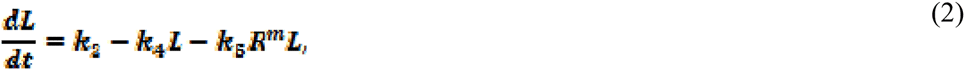

with suitable initial conditions *R*(0)=*R*_0_ and *L*(0)=*L*_0_. Here the function *f*(*t*) is given by (1+*c*)*k*_1_at times *t* ∈ [*t_on_,t_off_*]when IR is inflicted on the cell, and it is given by *k*_1_ otherwise. In this model, the influence of IR on the system is represented through the time-dependent function *f*(*t*) that describes the rate ROS generation. Lysosomal permeability is represented through the loss term – *k*_*f*_ *R*^*m*^ *L* in equation (2), which causes the number of viable lysosomes to decrease as the number of ROS increases (due to generation by IR).

## Results

### X-ray exposure induces rapid DNA damage in HF19 cells

Our results show that IR in the form of X-rays is able to significantly induce DNA damage across a 24-hour time period post exposure. To assess the sensitivity of HF19 cells to IR we used the alkaline comet assay. HF19 cells exposed to 0.1 Gy showed significant induction of DNA damage over the first 24 hours post-irradiation, peaking at 30 minutes (Figure 3. 13.74% ± 0.54). After 1 hour the mean tail DNA had fallen to 3.84% ± 0.23 (Figure 3), indicating active DNA repair. Over the remaining 24 hours DNA was slightly elevated, until 24 hours, at which time there was no change compared to the control. The response after exposure to 2 Gy X-ray demonstrated a substantial increase in DNA damage, 30 minutes (Figure 3. 28% ± 0.59) post-irradiation. The quantity of damage over the remaining 24 hours gradually decreased, but remained significantly elevated above control.

**Figure 3:**
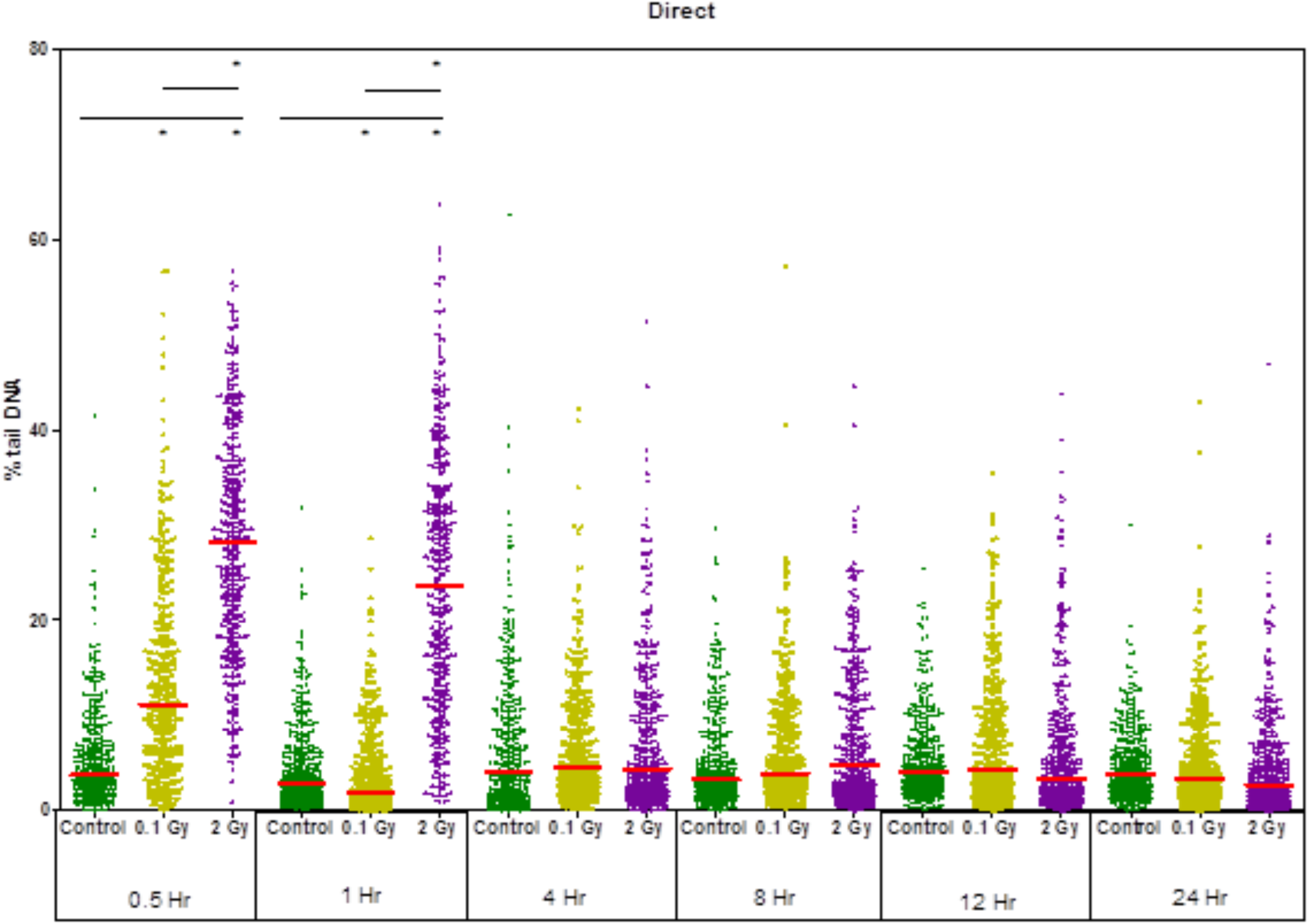
DNA damage was assessed in HF19 cells irradiated with 0, 0.1 and 2 Gy X-rays. Cells were assessed over a 24 hour period using the alkaline comet assay. IR induced significant levels of DNA damage; these were significantly elevated in 2 Gy irradiated samples until 8 hours. The amount of damage was dependent on dose (* p = < 0.05, error bars show SEM).

### X-ray exposure induces lysosomal damage up to 1 hour after exposure

IR was also able to induce significant changes in lysosomal permeability following X-ray exposure, particularly after 2 Gy. Thirty minutes post-irradiation with 0.1 Gy X-ray, there was a significant (4-fold) increase in green fluorescence compared to that observed in the control (Figure 4). It is unclear from these data whether this was as a result of complete rupture of individual lysosomes or due to a general increase in permeability across a number of lysosomes, although the magnitude of the increase would suggest complete rupture. Levels were almost 7-fold greater in cells exposed to 2 Gy X-rays compared to control groups (Figure 4). This persisted up to 1 hour post-irradiation in 2 Gy irradiated cells, using AO relocation.

**Figure 4:**
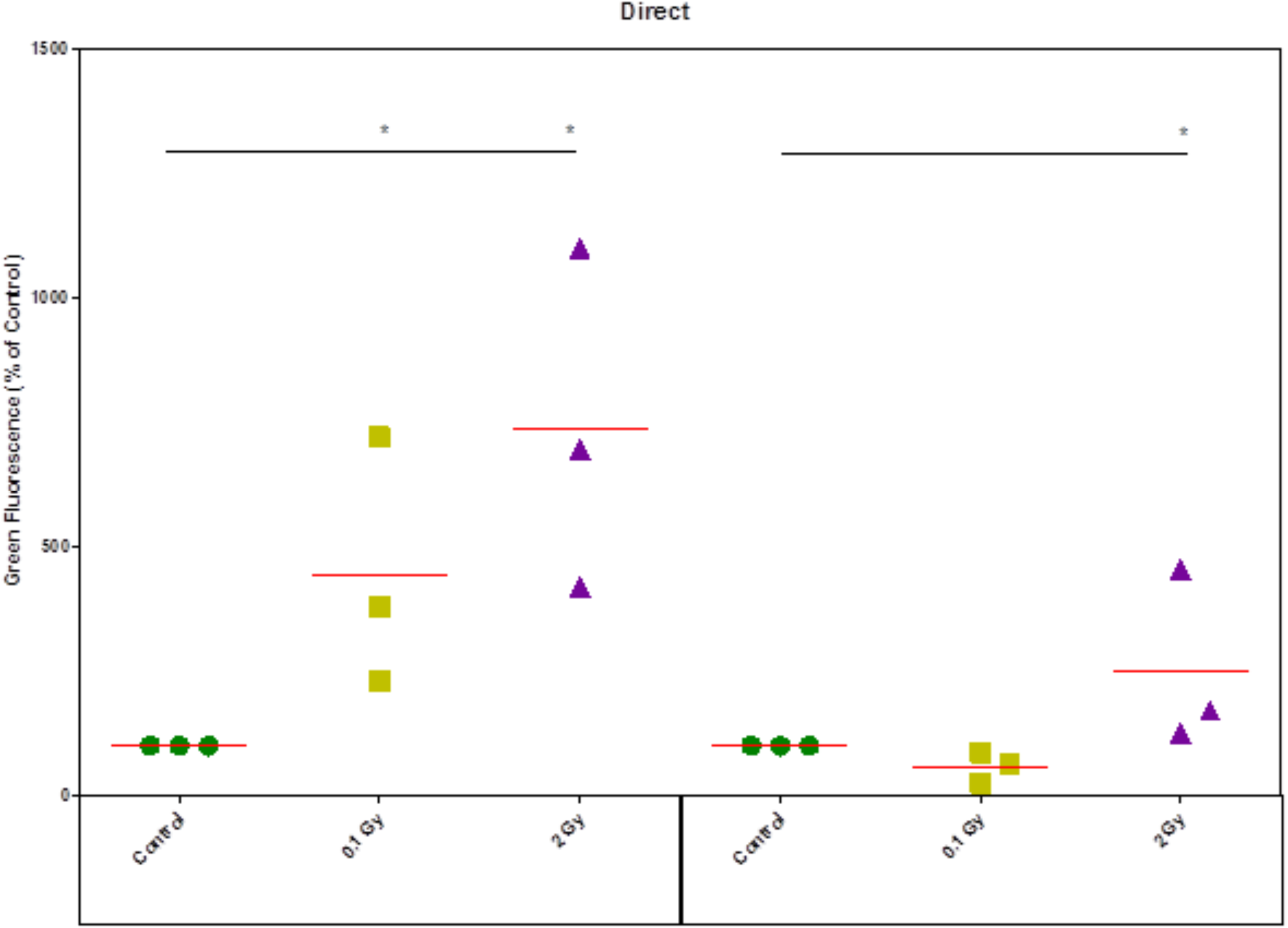
Lysosomal permeability was analysed using the acridine orange relocation method. HF19 cells were analysed 30 minutes and 1 hour post irradiation with 0, 0.1 and 2 Gy X-rays. Lysosomes showed significant levels of permeabilizatiom after 0.1 and 2 Gy irradiation, the effect persists to 1 hour in 2 Gy irradiated cells (* p = < 0.05, error bars show SEM).

### X-rays induce lysosomal permeability 4–24 hours post-irradiation

Lysosomal pertubations continued over the 24 hours following irradiation. A reduction in lysosomal number was observed at 24 hours for both doses. Initially we examined lysosomal number per cell, which was normalised to cell area. An oscilatory response was observed in cells that were exposed to 0.1 Gy over the first 12 hours while those that were exposed to 2 Gy were all increased above the control. We further explored the lysosomal profile following IR with respect to lysosomal fluorescence. In this instance both groups showed oscillatory responses to varying degrees (Figure 5), indicating subtle changes in permeability and possibly size.

**Figure 5:**
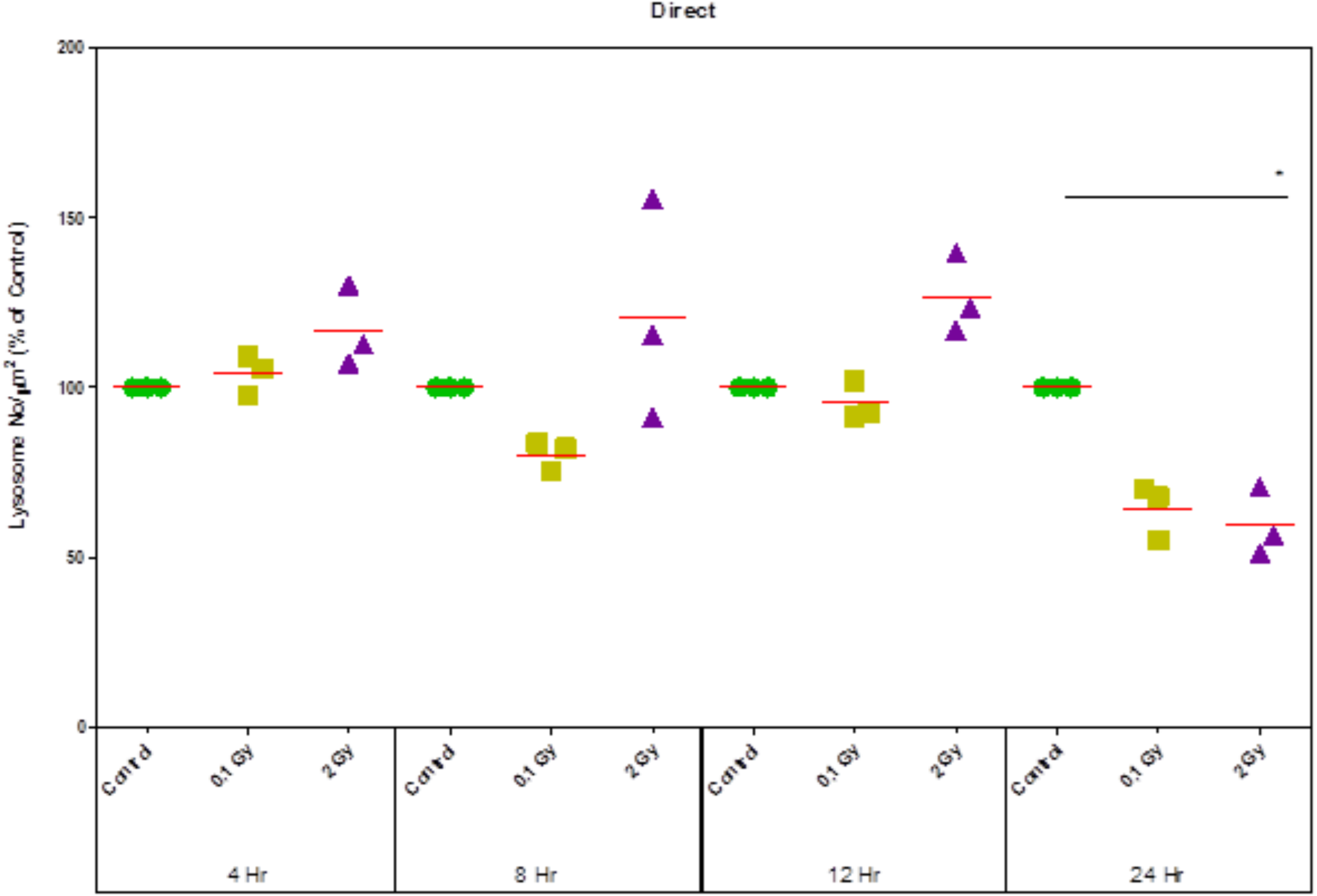
Lysosomal number was analysed using acridine orange and image analysis. HF19 cells were analysed from 4 hours to 24 hours post irradiation following exposure to 0, 0.1 and 2 Gy doses. Lysosome number fluctuated from 4 to 12 hours, however at 24 hours cells exposed to 2 Gy demonstrated a significant decrease in number. The effect was of a similar magnitude in cells exposed to 0.1 Gy, although this was not significant (* p = < 0.05, error bars show SEM).

### Persistent evelation of ROS up to 24 hours post-irradiation

ROS levels were elevated to a similar extent in both irradiated groups. This persisted over 24 hours. We analysed HF19 for levels of ROS using the general ROS marker, H_2_DCFDA. Thirty minutes post-irradiation, cells directly exposed to 0.1 Gy irradiation demonstrated a slightly elevated level of ROS (Figure 6. 125 % ± 2.10). ROS levels decreased further over the next 8 hours, but remained elevated above control samples. There appeared to a secondary increase in ROS at 12 hours post-irradiation (Figure 7. 119.39 % ± 2.77), followed by a reduction in ROS levels at 24 hours, but these levels remained approximately 10% above the control (Figure 7). 109.93 % ± 3.67).

**Figure 6:**
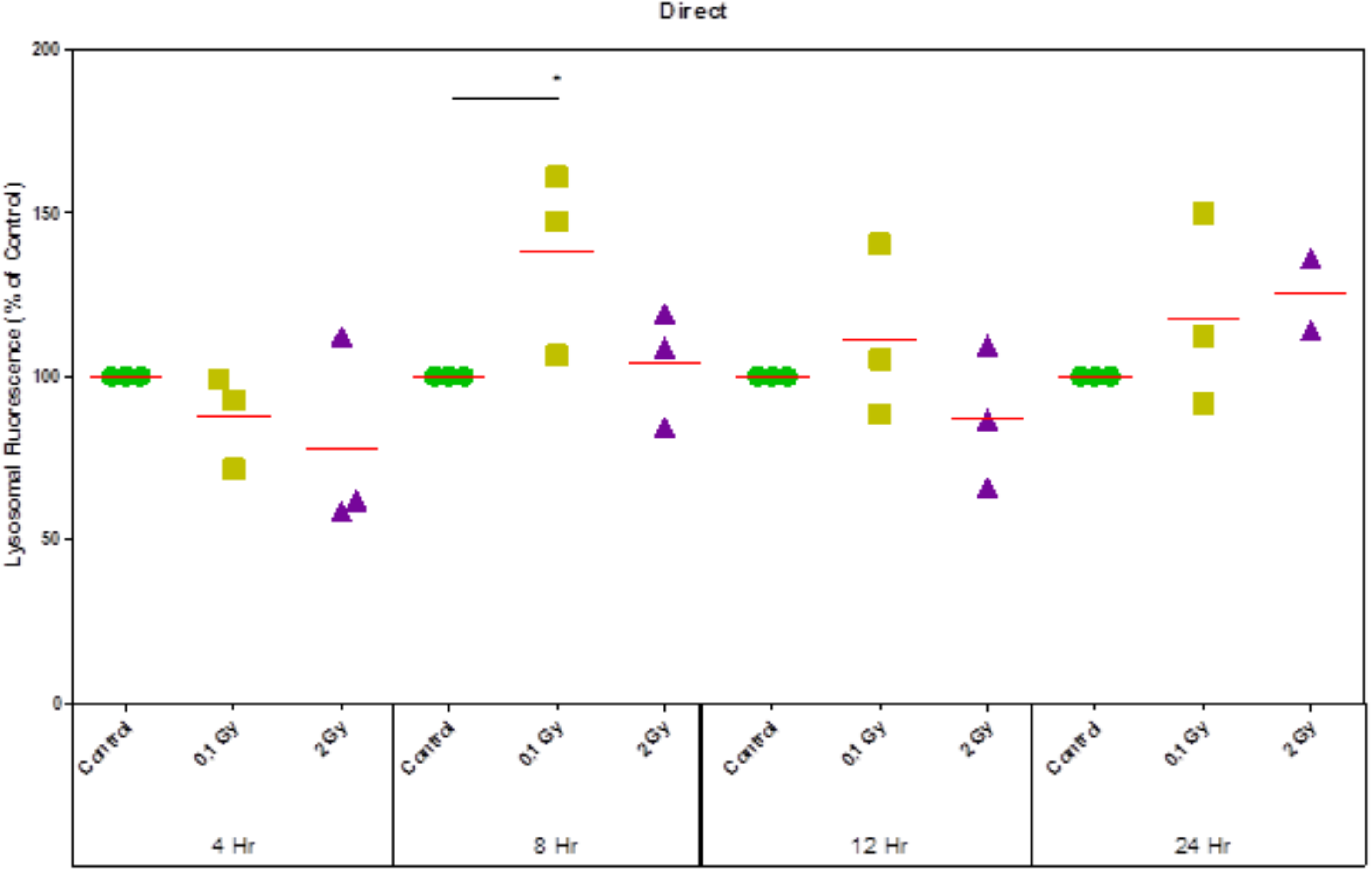
Individual lysosomal fluorescence was measured using the acridine orange uptake method. HF19 cells were analysed from 4 hours to 24 hours post irradiation following exposure to 0, 0.1 and 2 Gy doses. Fluorescence per lysosome was significantly increased 8 hours post 0.1 Gy exposure (* p = < 0.05, error bars show SEM).

**Figure 7:**
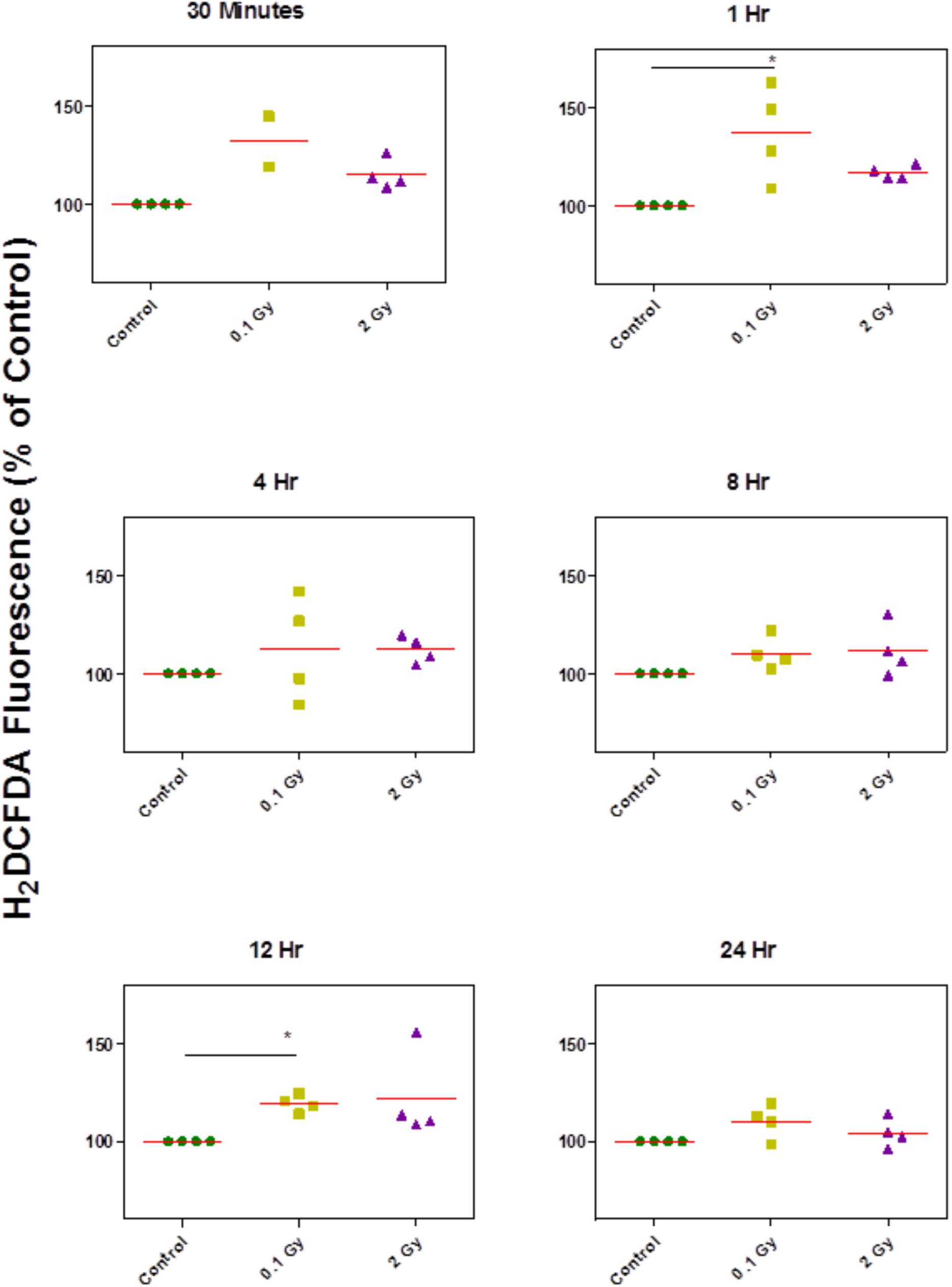
HF19 cells were analysed for oxidative stress using the H2DCFDA assay up to 24 hours post 0, 0.1 and 2 Gy X-ray irradiation. Levels were persistently elevated over the time course considered; the effect appeared to be independent of dose, as both 0.1 Gy and 2 Gy were very similar in level of increase (* p = < 0.05, error bars show SEM).

### Level of DNA damage following exposure to bystander signals

DNA damage was induced in bystander cells 1 hour after incubation with irradiated cell conditioned medium (ICCM). There were subtle changes in the effect, dependent on dose and time (Figure 8). The increase in damage was persistent in cells that received media from 2 Gy irradiated cells for the 24 hour time period. Cells that received media from 0.1 Gy irradiated cells showed similar levels of damage compared to the control at 8 and 24 hours but was elevated at all other time points.

**Figure 8:**
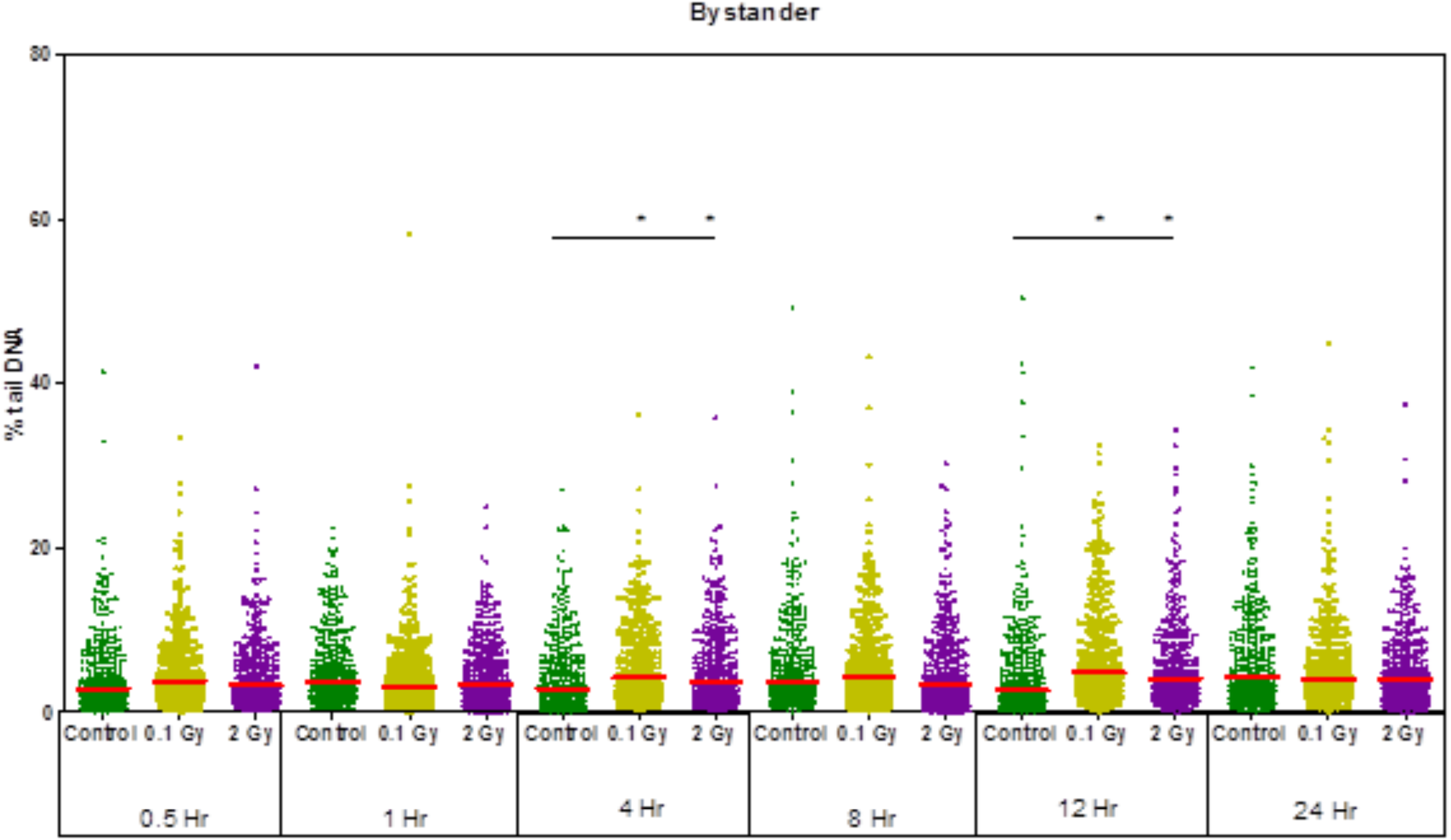
DNA damage was assesed in HF19 cells exposed to irradiated cell conditioned media (ICCM). Cells were asessed over a 24-hour period using the alkaline comet assay. ICCM was able to inuduce significant levels of DNA damage 1 hour after incubation with ICCM. Cells that received 2 Gy media showed persistent levels of DNA damage until 24 hours, whereas cells that recived ICCM from 0.1 Gy irradiated cells showed fluctuating levels of damage (* p = < 0.05, error bars show SEM).

### Lysosomal changes following bystander signals 30 minutres and 1 hour post-irradiation

Lysosomal stability was increased in both ICCM groups when compared to the control, as observed by a reduction in cellular green fluorescence (Figure 9). However, after 1 hour, lysosomal stability showed no significant difference. This was the same for both doses.

**Figure 9:**
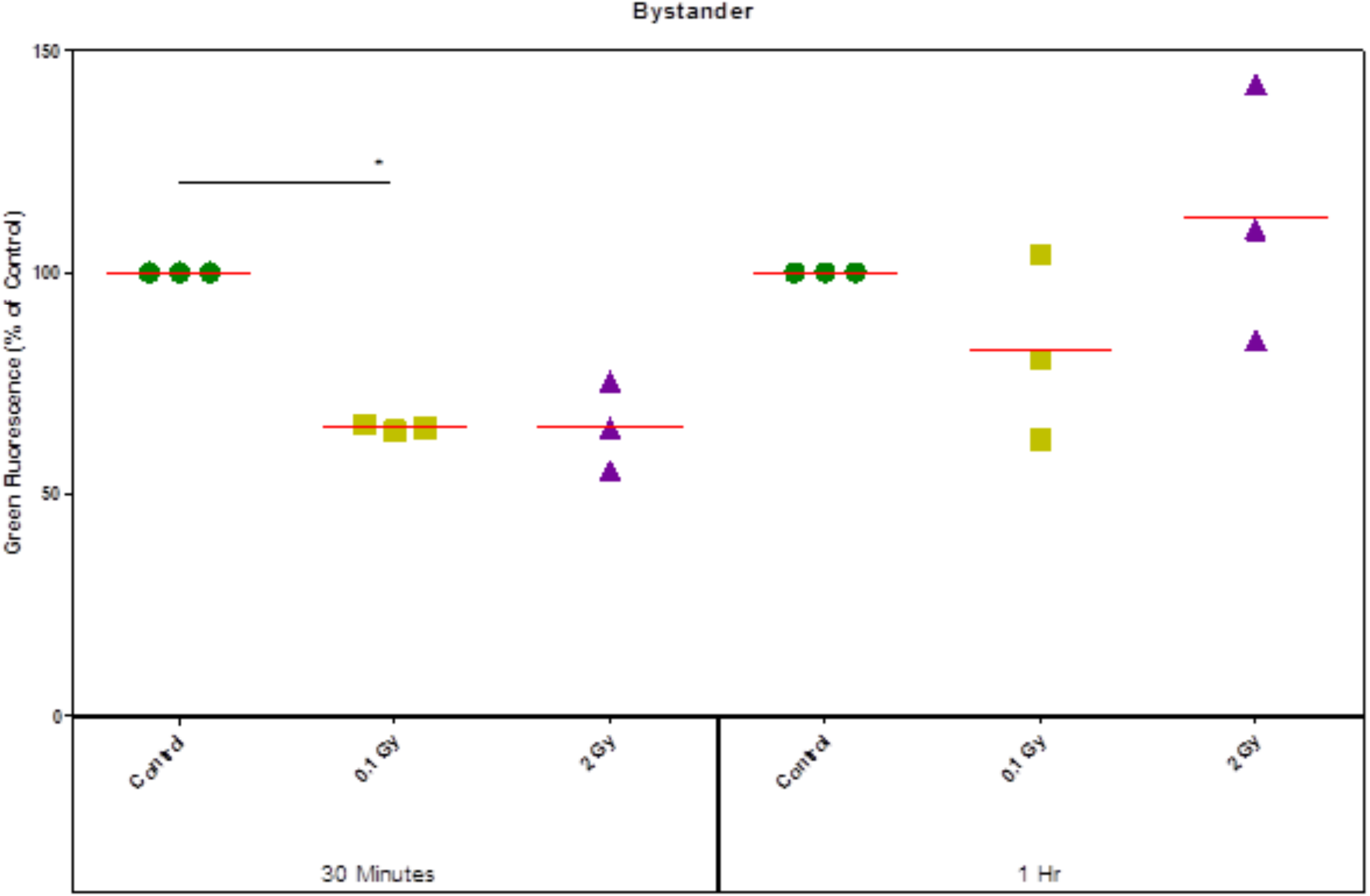
Lysosomal permeability was analysed using the acridine oranage relacation method. HF19 cells were analysed 30 minutes and 1 hour post incubation with media from 0, 0.1 and 2 Gy X-ray irradiated HF19 cells. Incubation with ICCM showed a reduction in fluoresence associated permeability, suggesting an increase in lysosomal volume or cell permeability. However after 1 hour there was no significant difference (* p = < 0.05, error bars show SEM).

### Bystander signals are able to induce lysosomal changes up to 24 hours post irradiation

ICCM was able to induce lysosomal alterations in the first 24 hours post incubation. Bystander cells also showed different effects from those seen in directly irradiated cells. Fours hours after incubation with 0.1 Gy ICCM there was a significant reduction in number of lysosomes however cells that that received 2 Gy ICCM demonstrated no significant change (Figure 10). The number proceded to fluctuate until 24 hours post incubation, at which point both ICCM groups showed an increase in number. This effect was greater in the cells that received 2 Gy ICCM (Figure 9).

**Figure 10:**
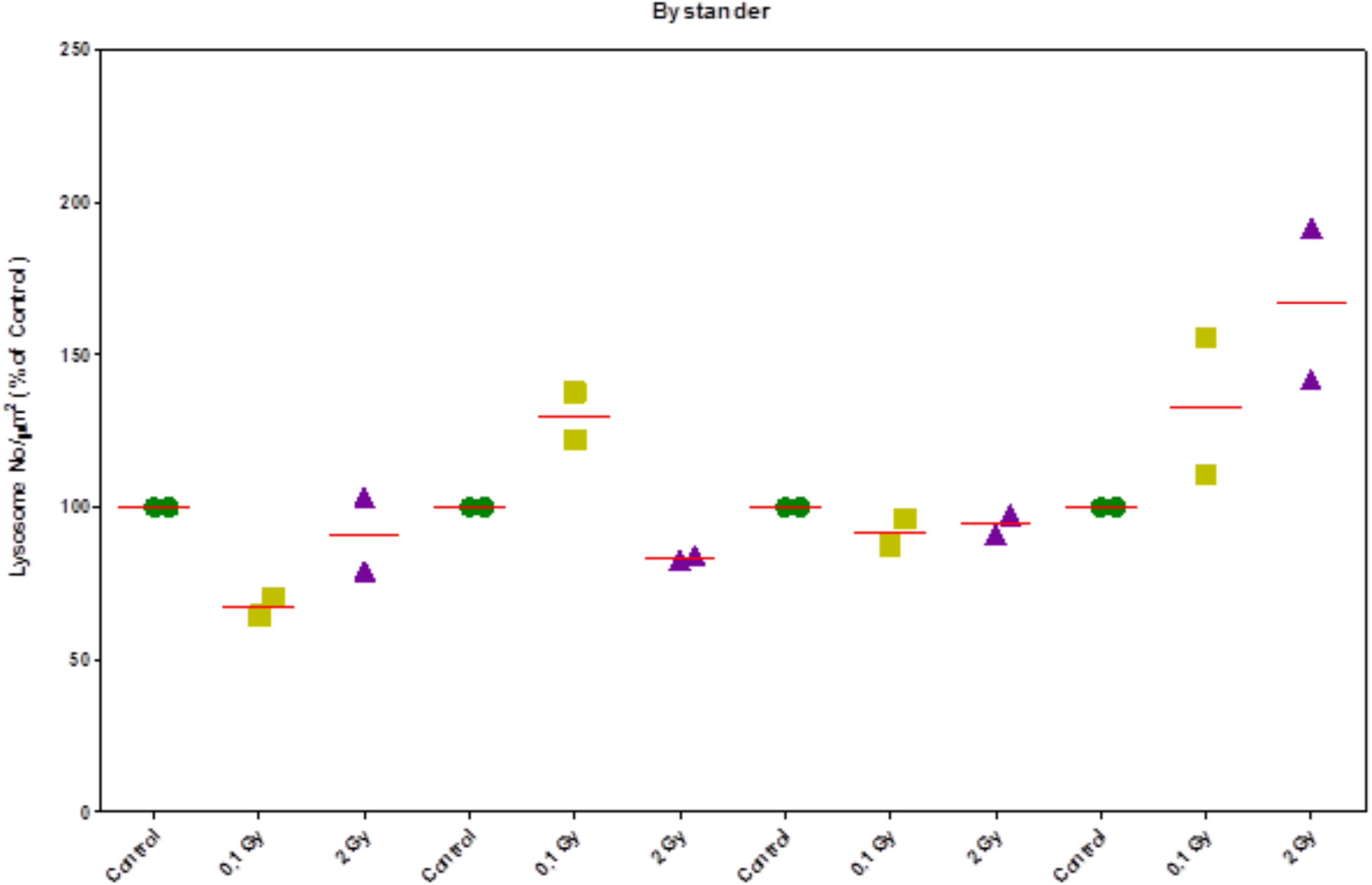
Lysosomal number was analysed using acridine orange and image analysis. HF19 cells were analysed from 4 hours to 24 hours post incubation with media from 0, 0.1 and 2 Gy X-ray irradiated HF19 cells. Numbers showed an oscillatory pattern; HF19 that received 0. 1 Gy ICCM showed significant reduction at 4 hours. At 24 hours, both doses showed an elevated number compared to the control bystander group (* p = < 0.05, error bars show SEM).

We also examined individual lysosomal fluoresence in order to ascertain if lysosomes may not be completely lysed but maybe more permeable. There was no signifcant change in ICCM groups 4 hours post incubation. However, there was a a clear and significant decrease in fluorescence per lysosome (Figure 11).

**Figure 11:**
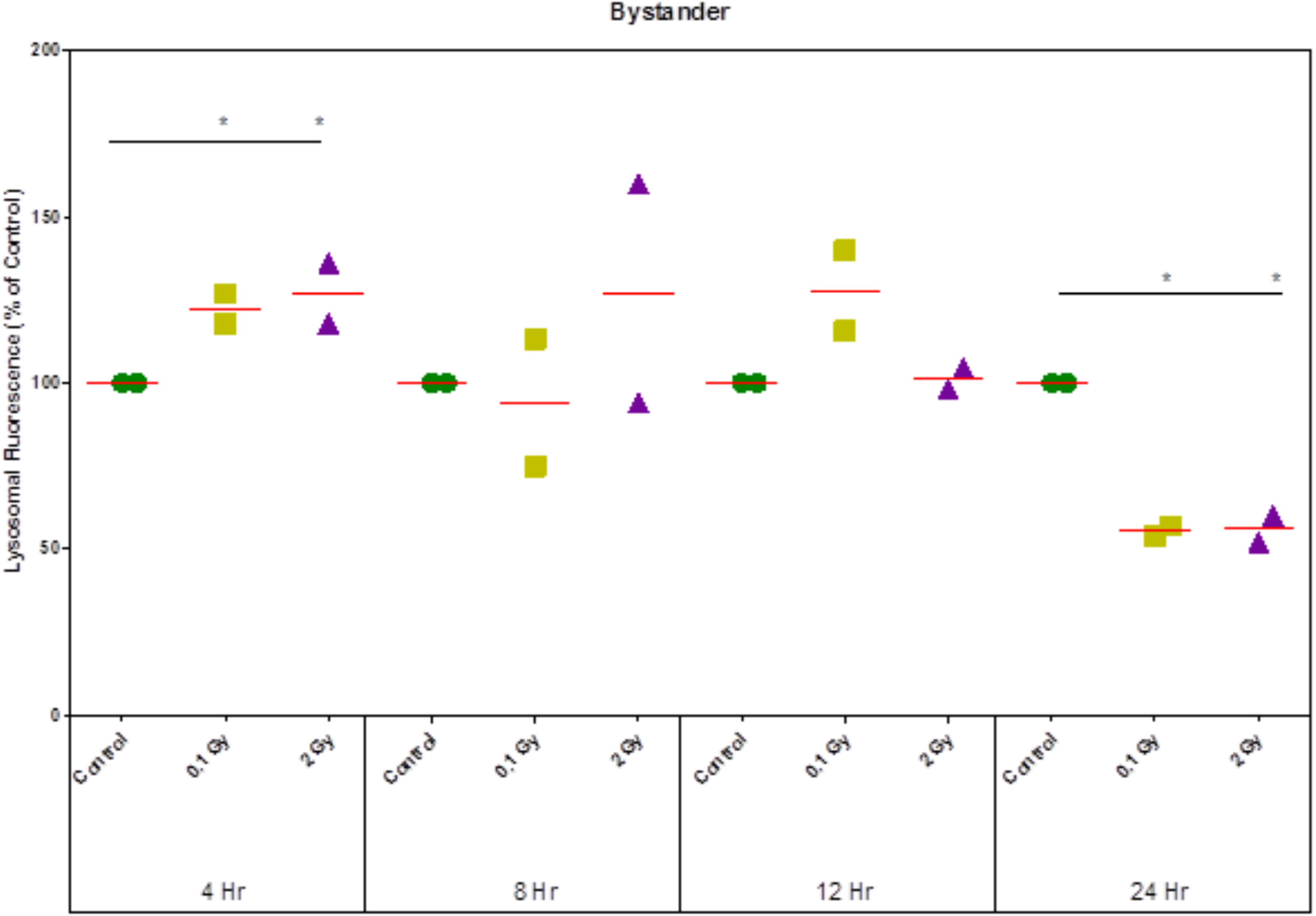
Individual lysosomal fluorescence was measured using the acridine orange uptake method. HF19 cells were analysed from 4 hours to 24 hours post incubation with media from 0, 0.1 and 2 Gy X-ray irradiated HF19 cells. Whilst lysosomal numbers did not change, there were changes in lysosomal fluorescence, indicating varying permeability. At 4 hours there was an elevation in fluorescence, indicating a more stable state. At 24 hours both ICCM bystander groups showed a reduction in fluorescence to a similar level, suggesting a more permeable state (* p = < 0.05, error bars show SEM).

### Bystander signals can alter oxidative stress

Bystander signals produced by irradiated cells had no effect on oxidative stress over the first hour post-irradiation (Figure 12). However, after 4 hours, oxidative stress was reduced. This effect was similar for both doses. Over the remaining time course, cells exposed to 0.1 Gy irradiated media showed no difference compared to the control (Figure 12), whereas cells exposed to 2 Gy bystander media showed small reductions in oxidative stress over the remaining time points, although this effect was not significant.

**Figure 12:**
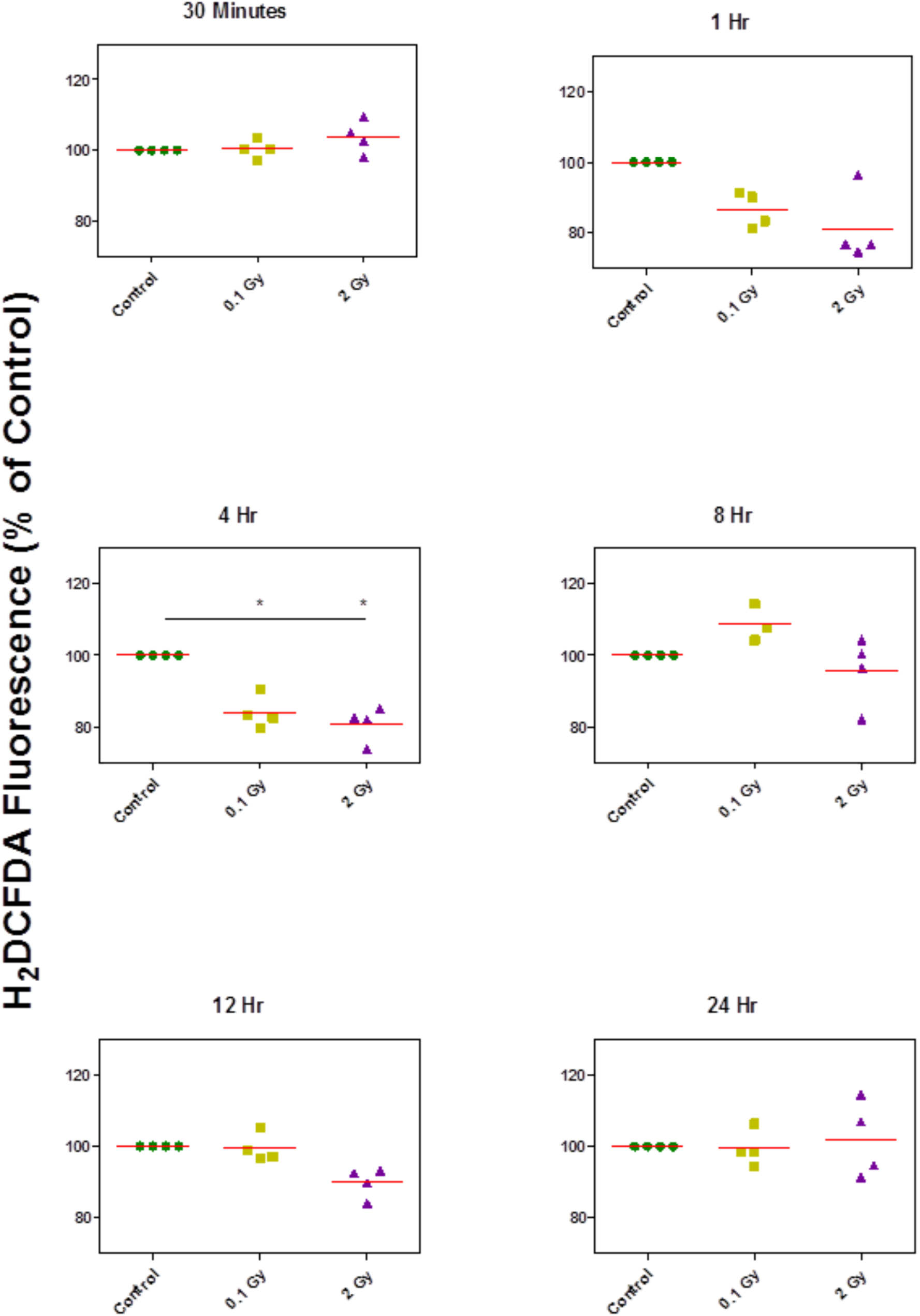
HF19 cells were analysed for oxidative stress using the H2DCFDA assay up to 24 hours post incubation with media from 0, 0.1 and 2 Gy X-ray irradiated HF19 cells. Although no effect was seen 30 minutes after incubation, the following time points showed a reduction in oxidative stress, which became significant at 4 hours. After 24 hours, levels had returned to that of the control (* p = < 0.05, error bars show SEM).

### Modelling suggests a possible mechanism for ROS-mediated persistent lysosomal permeability

In order to understand the experimental observations reported above, we now turn to our mathematical model of ROS-mediated lysosomal damage defined by equations (1)-(2). This model seeks to address hypothesised effect(s) of IR on ROS dynamics, and hypothesised interaction(s) between ROS and lysosomes, that could explain the observed dynamics post-IR, in which lysosomal damage recovery occurs over a relatively short timescale (less than 4 hours) while the levels of IR-induced ROS remain elevated for up to 24 hours.

Our first step is to seek steady state solutions (*R*\*,*L*\*)in the absence of IR, thus setting *f*(*t*) =*k*_1_. At steady state, setting equation (2) to zero yields

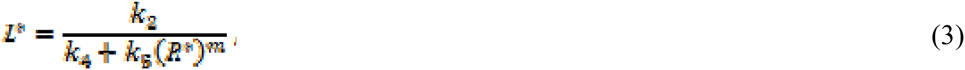

which may be substituted into (1) and rearranged to give the following quadratic equation that *R*\* must satisfy:

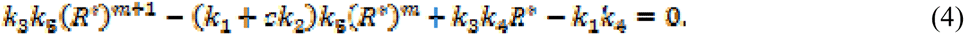

The number of distinct positive roots to equation (4), which gives the number of different biologically relevant steady states, may be found using Descartes’ rule of signs. This rule states that the number of positive roots to (4) is either equal to the number of sign differences between consecutive non-zero coefficients, or less than it by an even number. Now, as all rate constants are positive, there are three sign changes, hence either a single positive root or three positive roots. This means that the model either has a unique positive steady state, or three positive steady states. The latter case may correspond to bistability, in which a system evolves to one of two possible equilibria depending on its starting point. By exploring the space of model parameters, we identified a region of parameter space for which the system defined by equations (1)-(2) is indeed bistable. To illustrate this phenomenon, in Figure 13(a) we present a simulated time series in which a low dose of IR (corresponding to a small value of) is given to a cell for which the ROS and lysosome levels are at the ‘low’ steady state. We observed that the ROS and lysosome levels rapidly equilibriates back to their original values. However, as shown in Figure 13(b), when a high dose of IR (corresponding to a small value of) is given, the ROS level rapidly rises and reaches a distinct, ‘high’ steady state value. We explore the possible implications of this behaviour in our concluding discussion.

**Figure 13:**
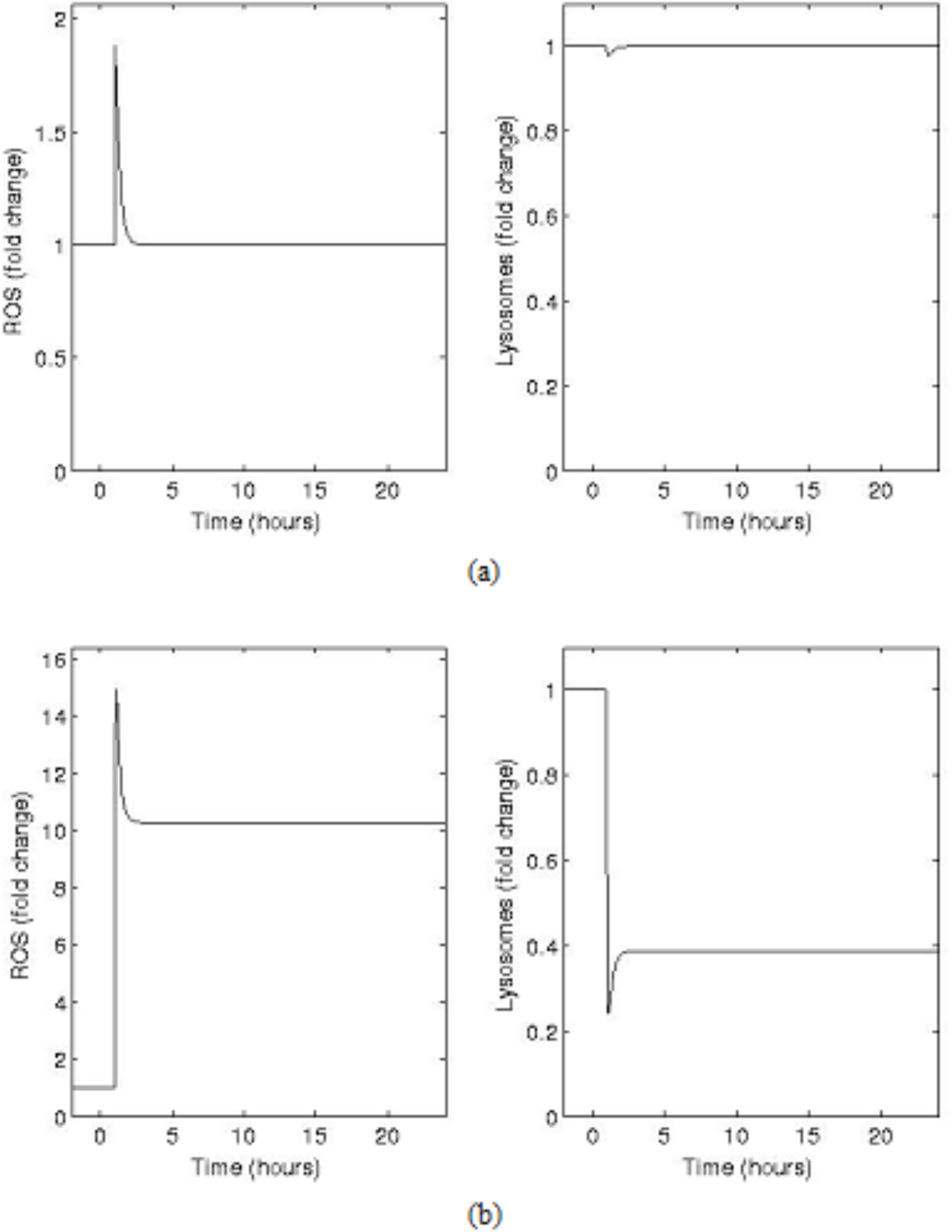
Simulation of the model of ROS-lysosome interactions defined by equations (1)-(2) in the case of ‘high’ and ‘low’ doses of IR. Parameter values are k1 = 2, k2 = 1, k3 = 7, k4 = 9, k5 = 1, m = 2, n = 40, ton = 1, ton = 1.0167, r0 = 0 and l0 = 0.1094. The time series for and are scaled by their initial value in each case. In (a), we take the IR damage ‘scale factor’, corresponding to a ‘low’ dose. In (b) we take, corresponding to a ‘high’ dose. In each case the numerical solution is obtained using a fourth-order Runge-Kutta method, implemented in MATLAB, with a sufficiently small upper bound placed on the time step to allow the window of IR damage to be resolved.

## Discussion

It is increasingly recognised that IR-induced effects are not limited to the induction of nuclear DNA damage, but that a number of intracellular targets, such as lysosomes, can also be affected after exposure. The lysosome contains enzymes capable of catalysing the breakdown of intracellular material including DNA, lipids and protein. It has been suggested that lysosomal DNase IIα released after lysosomal rupture can induce chromosomal aberrations much like those seen in IR-induced chromosomal instability *(19)*. Hence, the lysosomal response to IR is of critical importance to understand. The ability of IR to cause lysosomal damage may originate in a direct physical interaction with the lysosomal membrane; alternatively IR may act through an indirect mechanism, such as lipid peroxidation through radiation-induced ROS. Cell signalling through cytokines may also be involved *(18)*.

As expected ionizing radiation induced DNA damage in directly irradiated cells most likely through the production of ROS. DNA damage was also observed in bystander groups at 4 and 12 hours post irradiation. However no increase in oxidative stress was noted in the bystander groups, we therefore suggest the damage is likely attributable to inflammatory signals such as cytokine signalling *(39, 40)*.

We hypothesized that radiation, through a direct or indirect mechanism, was able to alter the lysosomal membrane and induce a state of increased permeability. Our results demonstrate that IR can alter lysosomal stability, not only in directly irradiated cells, but also in bystander populations. Our findings suggest that IR is able to induce significant lysosomal membrane permeabilization 30 minutes after exposure to 2 Gy X-rays, although the effect was also seen with a dose as low as 0.1 Gy. The induction of lysosomal membrane permeabilization was also observed 1 hour following exposure, although this was specific to 2 Gy exposure. Taken together, these results suggest that the process of lysosomal membrane permeabilization is not limited to a direct interaction, since it persists over an extended timescale. In fact, lysosomal changes are still detectable 24 hours after exposure to IR.

Changes in the homeostatic norm, such as aberrant ROS levels and associated downstream effects such as lysosomal lipid peroxidation, may account for the persistent changes observed in membrane permeability. A number of studies have linked ROS to lysosomal membrane permeabilization, although these have not considered the effects of IR *(41-43)*. Common features amongst these studies have linked the process directly to Fenton-type chemistry involving the transition metal iron. Iron is commonly found in the lysosome *(42)*. However, when iron encounters H_2_O_2_ it reacts to form other radicals, such as the hydroxyl and superoxide radicals. These radicals then interact with the membrane of the lysosome, initiating a chain reaction within the lipid membrane content inducing permeabilization *(44)*. This may be driving the early lysosomal membrane permeabilization following IR that was observed.

Bystander exposed cells demonstrated what appears to be an increase in lysosome number due to the reduction in cytoplasmic green fluorescence or alternatively an increase in the cell membrane permeability, perhaps as a result of the bystander signal. This observation occurred in conjunction with a small but statistically significant increase in DNA damage. The bystander signal therefore was not only able to change lysosomal/cell membrane properties but also DNA damage. However it is unclear if these effects are dependent on one another.

Over the remaining 24-hour period, bystander cells exhibited oscillations in lysosomal number in response to the addition of conditioned media. Saroya, Smith, Seymour and Mothersill [45] observed similar oscillations in zebra fish when examining calcium flux. Other oscillatory responses to bystander conditions were proposed by Lev Bar-Or *et al. (46)* when examining the p53-mdm2 feedback loop. These patterns of response, particularly in the p53 pathway, could be influential in driving the processes discussed in this study.

Having observed changes in lysosomal membrane permeability following IR, we further hypothesized that this is attributable to ROS. It is well known, and has been reproduced in the present study, that IR is a potent stimulator of ROS within the cell. Due to the short-lived nature of ROS, we might expect the majority to be generated and removed within the first few minutes post-irradiation *(2, 47)*. However, our results show that either some of these species have a longer half-life or are being produced as an indirect effect of radiation exposure on the time scale of hours post irradiation. The elevation in oxidative stress is maintained through the time course from 30 minutes to 12 hours. We also observed a small increase in both doses at 12 hours, although this does not correlate with any pattern seen in lysosomal disturbances.

The initial radiation insult is likely to consume a large proportion of the cell’s antioxidant capacity. This, in combination with an increased production of ROS, could induce a change in the cells oxidant status and ultimately alter the direction in which the cell is heading for example from a path of cell differentiation to one of cell proliferation. Building on previous work by *(47, 48)* have suggested that delayed ROS production may be attributable to p53 and its potent transcriptional activity, in this instance up-regulating redox related genes responsible for ROS induction and mitochondrial failure. The effects seen in the present study might be attributable to such a mechanism, involving changes in protein expression.

Bystander cells showed a reduction in oxidative stress when exposed to ICCM. This effect was only noted after 1 hour and 4 hours, suggesting that the bystander signal may transfer some antioxidant property over an intermediate timescale. However, DNA damage was also increased as a result of the bystander signal. This positive and negative BE has been documented previously and is reviewed by *(49)*.

To further explore the possible mechanism by which ROS mediate persistent lysosomal damage, we constructed a simple mathematical model for the temporal dynamics of the intracellular levels of ROS and lysosomes. A key assumption in the model is that several ROS molecules at once are required to induce permeability in each lysosome leads to a non-linear dependence of lysosome rupture on the ROS level. This assumption of cooperativity, coupled with amplification of ROS production by damaged lysosomes is shown to be capable of leading to bistable behaviour, whereby IR is able to shift the cell from one redox state to a new equilibrium if the radiation dose exceeds a threshold level. This may have profound implications for normal homeostatic control of cellular processes such as cell proliferation. Clearly this model represents an abstraction of reality, where we would expect the process of lysosomal rupture to be more gradual with progressive hits by ROS. A further source of complexity that we have neglected in our model is that, over longer timescales, we would expect processes such as autocatalytic extinction of ROS through NADH to lead to a gradual return of ROS levels to their ‘low’ or ‘normal’ steady-level. Nevertheless, our model hypothesises an intriguing possible mechanism where the cell’s natural resistance to excess ROS is overcome through saturation of defence mechanisms, or alternatively through an orchestrated mechanism to increase rate of ROS production. As a result cells can exist into two states of ROS concentration for which IR can act as a switch. The initial state corresponds to normal desired cellular function; however the secondary state, corresponding to higher ROS concentration, has the ability to cause intracellular damage and change cell function. A natural next step would be to consider the intracellular localisation of ROS and lysosomes following IR and resolve their spatio-temporal dynamics, which may highlight the likely stochastic nature of this process. A further natural extension of the model is to consider the behaviour of bystander, rather than directly irradiated, cells.

In conclusion, we have investigated the link between ionizing radiation exposure, oxidative stress and the sub-cellular response in particular lysosomal stability in primary human fibroblasts. We have demonstrated for the first time that IR alters lysosomal membrane permeability at medically relevant doses in primary human fibroblasts. These doses also induce persistent increase in oxidative stress in directly irradiated cells. However irradiated cells produce antioxidant signals that are functional in a bystander population. Taken together, our results demonstrate a potential role for IR-induced ROS in lysosomal membrane permeabilization.

Persistent alterations in lysosomal membrane have many wider implications for cellular and tissue health. Persistent leakage of enzymes has the potential to drastically alter the cell signalling environment, with small increases in permeability possibly driving cell proliferation. Complete rupture, however, could induce apoptosis or necrosis, which is a driver of inflammatory processes in a tissue microenvironment. We anticipate that such processes may be involved in NTE, and warrant further investigation, it also adds weight to the idea of the whole cell being sensitive to radiation and contributing to NTE.

## Author contributions

SB performed experiments and wrote the paper. AF produced model and wrote paper. DF contributed to model and wrote paper. MK planned experiments and wrote paper.

## Acknowledgements

SB was funded by an Oxford Brookes Nigel Groome Studentship. AF is funded by the EPSRC through grant EP/I017909/1 (www.2020science.net). The authors would like to thank Drs Mark Hill and James Thompson and Mr Luke Bird for radiation expertise.

